# HMGB1 mediates the development of tendinopathy due to mechanical overloading

**DOI:** 10.1101/751495

**Authors:** Guangyi Zhao, Jianying Zhang, Daibang Nie, Yiqin Zhou, Feng Li, Kentaro Onishi, James H-C. Wang

## Abstract

Mechanical overloading is a major cause of tendinopathy, but the underlying pathogenesis of tendinopathy is unclear. Here we report that high mobility group box1 (HMGB1) is released to the tendon extracellular matrix and initiates an inflammatory cascade in response to mechanical overloading in a mouse model. Moreover, administration of glycyrrhizin (GL), a naturally occurring triterpene and a specific inhibitor of HMGB1, the tendon’s inflammatory reactions. Also, while prolonged mechanical overloading in the form of long-term intensive treadmill running induces Achilles tendinopathy in mice, administration of GL completely blocks the tendinopathy development. Additionally, mechanical overloading of tendon cells *in vitro* induces HMGB1 release to the extracellular milieu, thereby eliciting inflammatory and catabolic responses as marked by increased production of prostaglandin E_2_ (PGE_2_) and matrix metalloproteinase-3 (MMP-3) in tendon cells. Application of GL abolishes the cellular inflammatory/catabolic responses. Collectively, these findings point to HMGB1 as a key molecule that is responsible for the induction of tendinopathy due to mechanical overloading placed on the tendon.

## Introduction

Tendinopathy, a debilitating chronic tendon disorder, is manifested in clinical settings by a combination of pain, swelling, compromised tendon structure, and rupture (1). Tendinopathy, which involves tendon inflammation and degeneration, affects healthy individuals during their active and productive years of life resulting in tremendous healthcare costs and economic impact due to work-loss (2, 3). In particular, insertional tendinopathy, which is common in young athletes, often occurs in the tendon proper proximal to the insertion into the heel bone and accounts for about 20% of Achilles tendon disorders (4). It is well established that while normal physiological loading is essential for tendon homeostasis, mechanical overloading induces the development of tendinopathy, characterized by disorganized matrix, reduced numbers and rounding of tendon cells, fibrocartilaginous change, and neovascularization (5, 6). A current concept on the mechanisms of tendinopathy is that repetitive loading may lead to a mechanobiological over-stimulation of tendon cells resulting in an imbalance between the synthesis and breakdown of matrix proteins, especially collagen (7–9). The resulting mismatch is a continuous loss of collagen in the tendon by repetitive loading with insufficient recovery time, which initiates a catabolic degenerative response that leads to tendinopathy (6, 10).

It is now recognized that inflammation is part of tendinopathy and could lead to tendon degeneration that occurs at late stages of tendinopathy (11–13). Under excessive mechanical loading, abnormal levels of proinflammatory mediators may be released triggering inflammatory reactions and resulting in severe pain in tendon. PGE_2_, an enzymatic product of cyclooxygenase-2 (COX-2), is an established potent lipid mediator of inflammation and pain in tendinopathy (14). Previously, we showed that COX-2 and PGE_2_ are produced at abnormally high levels in tendons and by tendon cells subjected to mechanical overloading (15–17). Such an abnormal increase in PGE_2_ levels plays an important part in tendon inflammation (15, 18), which can lead to tendon degeneration characterized by hypercellularity, angiogenesis, and abnormal arrangement of collagen fibers thus impairing the structure and function of tendons (17, 19, 20). Additionally, failure to regulate specific MMP activities in response to repeated mechanical loading accelerates tendon degeneration (21). Nevertheless, the identity of the molecular mediators through which mechanical overloading triggers the production of these inflammatory/catabolic mediators that eventually leads to tendinopathy remains largely unknown.

As a non-histone nuclear protein, HMGB1 is recognized as an endogenous danger signaling molecule that triggers inflammatory responses when released into the extracellular milieu. While HMGB1 is present in the nuclei of almost all cells where it regulates DNA stability and gene expression (22, 23), it can be released from a variety of cells especially macrophages as a result of an active process in live cells, or passively released from stressed, injured and necrotic cells (24). Indeed, mechanical loading *in vitro* or *in vivo* induces HMGB1 release from ligament cells to the extracellular matrix (ECM) and participates in the inflammation process and tissue remodeling by modifying the local microenvironment (25, 26). In short, once released, HMGB1 induces and maintains an inflammatory response (27–30). Extracellular HMGB1 acts as an inflammatory mediator and triggers an inflammation cascade inducing the production of IL-1β, IL-6, and TNF-α (31, 32). HMGB1 also maintains that response by inducing its own release from monocytes and macrophages (33). Extracellular HMGB1 plays a key pathogenic role in many major diseases such as cancer, stroke, endotoxemia, and joint disorders (34, 35). Although extensive literature is available on the inflammatory role of HMGB1 in fibrosis and other diseases (31), only limited studies have linked HMGB1 to tendinopathy. Millar *et al*. have suggested a new mechanistic role of alarmins such as HSP70 in initiating inflammation in early stage tendinopathy (36). They also indicated that HMGB1 may play a pivotal role in the pathogenesis of a variety of inflammatory conditions; they further conducted a clinical study and found high levels of HMGB1 in tendinous tissues of supraspinatus tendinopathy patients (37). Increased expression of alarmins including HMGB1 has also been reported in another study with a few supraspinatus tendinopathy patients (38). Recently, upregulation of HMGB1 has been associated with inflammatory responses and ECM disorganization in rat rotator cuff tendon injury model (39).

However, whether HMGB1 mediates the development of tendinopathy due to mechanical overloading placed on the tendon is largely unknown. To determine this, we performed mouse treadmill running experiments. We report that in response to such mechanical overloading *in vivo*, HMGB1 was released to tendon matrix and initiated an inflammatory cascade, and this inflammation was inhibited by administration of glycyrrhizin (GL), a naturally occurring triterpene and a specific inhibitor of HMGB1. Furthermore, mechanical overloading in the form of long-term intensive treadmill running induced Achilles tendinopathy in mice, and administration of GL completely blocked the tendinopathy development. A detailed report is as follows.

## Materials and methods

### Ethics Statement

All experiments were performed in accordance with relevant guidelines and regulations. All animal experiments were approved by the Institutional Animal Care and Use Committee of University of Pittsburgh (IACUC protocol #17019968).

### Mouse treadmill running experiments

#### 1) Short term treadmill running experiments with various intensities

In total, 48 female C57B6/L mice (3 months old) were used for the *in vivo* treadmill running experiments and were divided into 4 groups with 12 mice in each group. The control mice remained in cages and were allowed cage activities. The remaining three groups ran on the horizontal treadmill but at different intensities; i) moderate treadmill running (MTR), ii) intensive treadmill running (ITR), and iii) one-time treadmill running (OTR). The running speed for all regimens was 15 meters/min. In the first week, mice were trained for 15 min to accommodate them to the treadmill running protocol and environment. In the following 3 weeks, mice in the MTR group ran for 50 min and those in the ITR group ran for 3 hrs a day, 5 days a week. Mice in the OTR ran for more than 5 hrs until fatigue. Performance of the mice (i.e., running time) was recorded to recommend inclusive/exclusive criteria. Immediately after running, the Achilles and patellar tendons were harvested from four groups of mice. Half of the tendon samples were used for ELISA and the remaining half was used for immunostaining.

#### 2) Short term intensive treadmill running (ITR) with GL administration

In these experiments, we used a total of 24 female C57B6/L mice (3 months old) with 6 mice in each of the 4 groups; i) cage control group (Cont) where mice did not receive any treatment and served as control group with intact tendon, ii) GL injection only where mice received daily intraperitoneal (IP) injection of GL (50 mg/kg body weight, Cat # 50531, Sigma-Aldrich, St. Louis, MO), iii) ITR group where mice ran on the ITR regimen (see above *in vivo mouse treadmill running model* for details), and iv) ITR with GL injection (GL+ITR) group where mice received daily IP injection of GL 15 min before the beginning of ITR regimen. The dosage of GL was selected based on previous studies (40–42). After treadmill running for 3 weeks, patellar and Achilles tendons were dissected out and the right and left side of each tendon from a single mouse were homogenized in T-PER buffer (Cat # 78510, ThermoFisher, Pittsburgh, PA) and the supernatants were used for ELISA to measure PGE_2_ and MMP-3.

#### 3) Long term ITR (Lt-ITR) with GL administration

This treadmill running protocol was similar to the 3-week running protocol, using a total of 24 mice divided into 4 groups. The only difference was that the Lt-ITR mice and Lt-ITR+GL mice ran a horizontal treadmill in the first 12 weeks, and then ran a 5° uphill treadmill for additional 12 weeks to increase the load on Achilles tendon to maximize the treadmill running effect. At the end of 24 weeks, all mice were sacrificed, and the Achilles tendons were dissected and used for histological and immunohistochemical (IHC) analyses.

### HMGB1-alginate beads implantation in tendon *in vivo*

To assess the function of HMGB1 *in vivo*, we developed a system to deliver HMGB1 into tendons *in vivo* to mimic long term release of HMGB1 induced by repetitive mechanical loading. Our delivery system consisted of a degradable polymer called alginate that contained HMGB1 to ensure local and continuous delivery of HMGB1 to maximize the effect in a relatively short period of time.

A 2% alginate solution was first prepared by dissolving alginate powder (Cat # 180947-100G, Sigma-Aldrich) in double distilled water with vigorous vortexing. Then, HMGB1 powder (Cat # H4652, Sigma-Aldrich) was added to the 2% alginate solution at the concentration of 0.5 mg/ml. Using a pipette, about 5 µl of the HMGB1-alginate solution was then added to 2 mM CaCl_2_ solution in the form of drops, which solidified to form alginate beads. Control alginate beads were prepared without adding HMGB1. The beads were then removed from the CaCl_2_ solution and allowed to air dry. The final diameter of the beads was around 0.5 mm, which is about 1/6 of the rat patellar tendon width. This protocol was developed in our laboratory (43).

Sprague Dawley (SD) rats (female, 6 months) were sedated by inhaling 2-3% isoflurane. The skin over the patellar tendon was then shaved, sterilized and a small incision was made on the skin to expose the tendon. HMGB1-alginate beads containing 2.5 µg HMGB1 or blank alginate beads with the same size for control were implanted into the central part of the left and right patellar tendons. After 2 and 4 weeks, 5 rats in each group were used for hematoxylin & eosin (H&E) and IHC staining to evaluate structure and compositional change in the patellar tendon tissue.

### ELISA for measuring HMGB1, PGE_2_, and MMP-3 in tendon

ELISA kits were used to measure HMGB1, PGE_2_, and MMP-3 protein levels in tendinous tissues. Briefly, mice Achilles and patellar tendinous tissues were weighed and blunt separated with forceps and soaked in 200 µl PBS for 24 hrs at 4℃ to allow HMGB1 in the matrix to diffuse to PBS. This was done to prevent nuclear HMGB1 leaking into the extracellular space thus allowing precise quantification of HMGB1 that was released to the extracellular space. The samples were centrifuged at 2,000 g for 30 min at 4°C and the supernatants were collected to measure HMGB1 concentrations using an ELISA kit (Cat # ST 51011, Shino-Test Corporation, Tokyo, Japan) according to the manufacturer’s instructions. All samples were analyzed in duplicates. To determine whether the above PBS extraction method collected HMGB1 from the tendon matrix only, we used an ELISA kit (Cat # ab156895, Abcam, Cambridge, MA) to measure the DNA concentrations in all samples lysed with 200 µl RIPA buffer (Cat # R0278, Sigma-Aldrich).

For PGE_2_ and MMP-3 measurements, the samples were vigorously homogenized with BioMasher Standard (Cat # 9790A, Takara, Shiga, Japan) in 200 µl T-PER tissue protein extraction reagent instead of PBS. The samples were then centrifuged as described above and the supernatants were collected for ELISA and the concentrations were determined using ELISA kits for PGE_2_ (Cat # 514010, Cayman, Ann Arbor, Michigan) and MMP-3 (Cat # LS-F5561, Lifespan Bio, Seattle, WA).

### Alcian blue and nuclear fast red staining

Alcian blue staining was performed using a kit (Cat # ab15066, Abcam) following the manufacturer’s protocol. Briefly, glass slides with tissue on it were hydrated first and incubated in acetic acid for 3 min. The slides were incubated in Alcian blue (1% solution, pH 1.0) solution for 30 min at room temperature and then rinsed with acetic acid. They were then rinsed with running tap water for 2 min, followed by washing with two changes of distilled water. The slides were stained with Alcian blue solution for an additional 5 min, followed by rinsing with running tap water and two changes of distilled water. The slides were counterstained, dehydrated with graded alcohols, washed in with xylene and covered with cover slips.

Alcian blue and nuclear fast red dual staining was performed as follows. The tissue sections were fixed with 4% paraformaldehyde for 20 min at room temperature, and then washed three times with PBS. The slides were stained with Alcian blue as described above, then washed with water 3 times and counterstained in 0.1% nuclear fast red solution (Cat # ab146372, Abcam) for 5 min. The slides were washed with water 3 times, and dehydrated through 95% alcohol and absolute alcohol, 3 min each. The slides were finally treated with xylene and mounted with resinous mounting medium. The photographs were taken with a histology microscope. With this staining, the nuclei appear as pink to red and glycoproteins as dark blue.

### Immunostaining of tendinous tissue

For immunostaining of tendinous tissue, Achilles and patellar tendons dissected from the mice were immediately immersed in O.C.T compound (Sakura Finetek USA Inc, Torrance, CA) in disposable molds and frozen at −80°C. Then, cryostat sectioning was performed at −25°C to obtain about 10 µm thick tissue sections, which were fixed in 4% paraformaldehyde for 15 min and blocked with universal blocking solution (Cat # 37515, ThermoFisher Scientific). The sections were then incubated with rabbit anti-mouse HMGB1 antibody (1 µg/ml, Cat # ab18256, Abcam) at 4°C overnight followed by goat anti-rabbit secondary antibody conjugated with Cy3 for 1hr at room temperature (0.5 µg/ml, Cat # AP132C, Millipore, Billerica, MA), and then counterstained nuclei with Hoechst 33342. Since the purpose of this staining was to evaluate the presence of HMGB1 in the extracellular milieu, the tissue sections were not treated with Triton X-100 to block the permeation through the nuclear membrane. HMGB1 levels in each tendon sample were normalized to the corresponding tissue weight.

For CD31 and CD68 staining, patellar tendons were harvested from 3 rats that received HMGB1-alginate bead or control bead implantation and tissue sections were prepared for immunostaining as described above. Anti-rat CD31 antibody (1 µg/ml, Cat # ab64543, Abcam) was used to detect endothelial cells and vessels and anti-CD68 antibody (1 µg/ml, Cat # ab955, Abcam) was used to detect monocytes/macrophages following the same procedure as above. H&E staining was used to evaluate overall tendon structure and cell density.

For collagen II (Col II) staining, the fixed tissue sections were treated with 0.05% trypsin for 20 min at 37°C and washed with PBS three times. Then, the washed tissue sections were reacted with rabbit anti-collagen II antibody (1:500, Cat. # ab34712, Abcam) at 4°C overnight. For SOX-9 staining, the tissue sections were further treated with 0.1% of Triton X-100 for 30 min at room temperature and washed with PBS another three times, then the sections were incubated with rabbit anti-SOX-9 (1:500, Cat # AB5535, Millipore) antibody at 4°C overnight. Finally, the tissue sections were washed 3 times with PBS and incubated with Cy3-conjugated goat anti-rabbit IgG antibody at room temperature for 2 hrs. Slides were then counterstained with Hoechst 33342.

### Statistical Analysis

Wherever appropriate, student’s *t*-test, or one-way ANOVA was used followed by Fisher’s least significant difference (LSD) test for multiple comparisons. When P-values were less than 0.05, the two groups compared were considered to be significantly different.

## Results

### Mechanical overloading *in vivo* induces HMGB1 release into tendon matrix

As the first goal of this study, we determined whether mechanical overloading in the form of mouse treadmill running induces the release of HMGB1 to the ECM *in vivo*. For this, first we determined the presence and localization of HMGB1 using Western blot and IHC in normal tendinous tissues without mechanical loading. The Western blot results showed that HMGB1 is present in the patellar and Achilles tendinous tissues, and immunofluorescence results showed that it is localized in the nuclei, not in ECM (**S1 Fig**). After treadmill running, Achilles tendon sections of the control group (cage activity only) showed the presence of tendon cells that stained blue with Hoechst 33342, but the tendon matrix was not positively stained indicating the absence of HMGB1 in the matrix (Fig 1A). The tendinous tissue was cut into 10 µm thickness pieces were not penetrated with detergent to minimize staining of HMGB1 in the nucleus (a penetrated tissue staining sample is shown in **S1C Fig**). Some peripheral areas that appear red are paratenon, which surrounds the tendon proper (Fig 1A, a, e). In the tendon sections from mice on the MTR regimen, HMGB1 staining was absent in the tendon matrix except for the mild positive staining in the peripheral areas (Fig 1A, b). However, a marked increase in HMGB1staining was observed in the mouse tendon matrix with the ITR regimen for 3 weeks (Fig 1A, c). A 20x magnification shows clear positive staining for HMGB1 outside the tendon cells and in the tendon matrix (arrowhead, Fig 1A, g). The same increasing trend for HMGB1 staining was observed in the tendon matrices of mice 5-7 hrs on the OTR regimen (Fig 1A, d, h). The majority of HMGB1 is detected surrounding the elongated tendon cells, which indicates HMGB1 is released by these cells.

**Fig 1.**
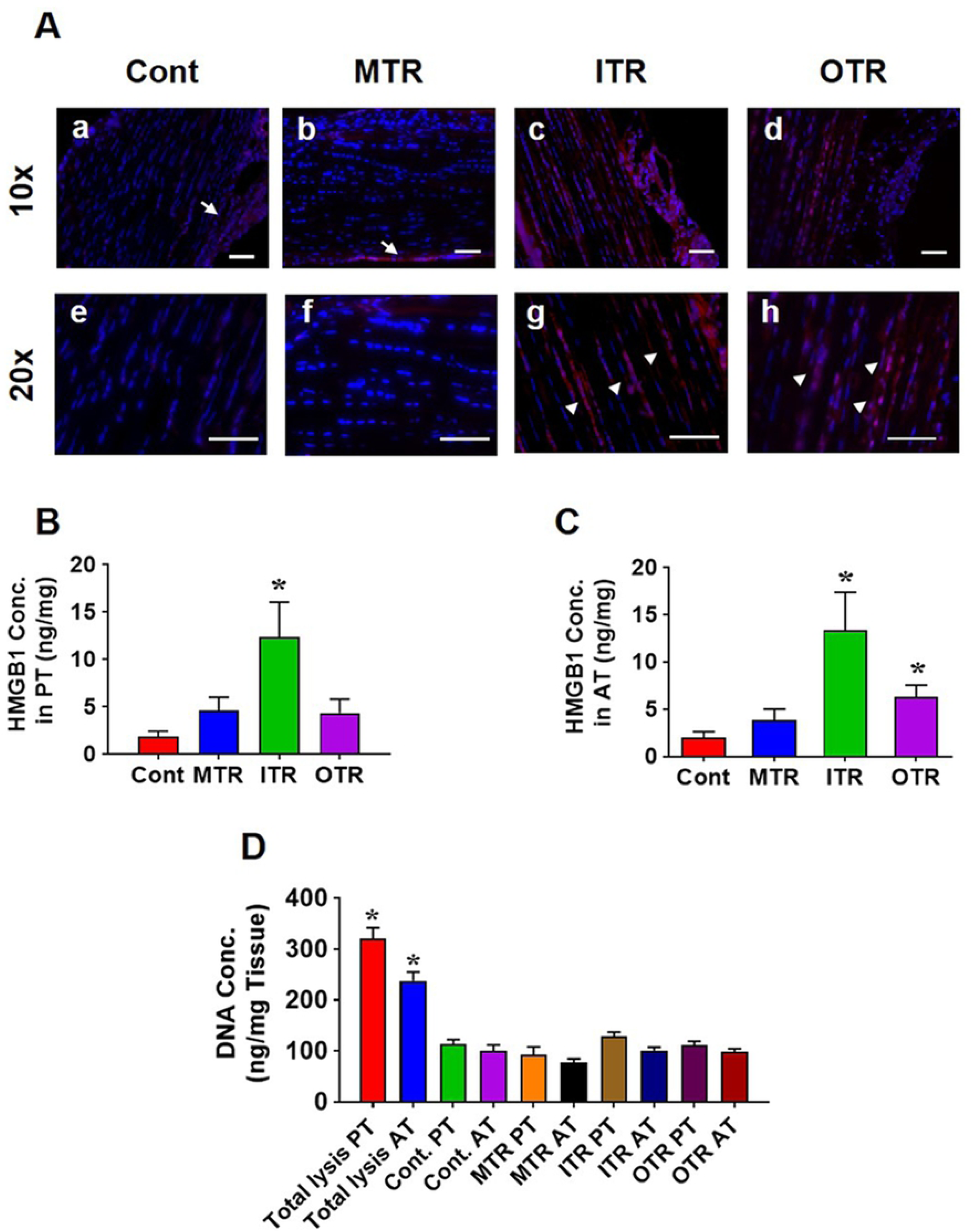
Mechanical overloading through mouse treadmill running increases HMGB1 levels in tendon matrix. (**A**) Immunostaining for HMGB1 under various mechanical loading conditions. (**a**) Achilles tendon (AT) from cage control mouse shows minimal HMGB1 staining in tendon matrix. (**e**) 20x magnification of (**a**) clearly shows the absence of HMGB1 staining in the matrix. (**b**) A representative tendon section from moderate treadmill running (MTR) group showing no positive stain for HMGB1 in the matrix. (**f**) 20x magnification of (**b**) showing negative stain for HMGB1. (**c**) Tendon matrix shows strong positive stain in the intense treadmill running (ITR) group indicating that HMGB1 has released to the matrix. (**g**) 20x magnification of **c** clearly shows positive stain in the matrix. Arrows point to positive staining. (**d**) Similar HMGB1 positive staining in the matrix of tendon section from one-time treadmill running (OTR). (**h**) 20x image of **d** (arrowheads point to positive staining). Note that the sections were not permeabilized with detergent to avoid the staining of HMGB1 in the nuclei. Also, the pink stains observed in the periphery of **a** and **b** (arrows) are paratenon or adjacent connective tissue. Data shown are the representatives from two independent experiments (n = 6 mice in each group). (**B**) ITR significantly increases HMGB1 levels, but OTR and MTR do not significantly alter HMGB1 levels in patellar tendons (PT) compared to control. (**C)** ITR and OTR significantly increase HMGB1 levels in Achilles tendons (AT) compared to control. There is no significant change in MTR group. HMGB1 measurement is normalized to tissue weight. (**D**) DNA concentrations of tendon samples are equivalent and significantly lower than total lysed sample. This means that the high HMGB1 concentrations in ITR and OTR are not due to excessive disruption of cells since HMGB1 releases together with DNA while cells are disrupted during tissue procession. Data represent mean ± SD. n = 6. \**P* < 0.05. Bar: 50 µm.

These findings were also confirmed by ELISA measurement of *in vivo* HMGB1 levels in mouse patellar and Achilles tendon matrices (PT and AT, respectively) subjected to mechanical loading protocols (Fig 1B, C). Specifically, HMGB1 levels were 6.6-fold higher in Achilles tendon, and 6.8 times higher in patellar tendon of ITR group when compared to the control mice that remained in cages. HMGB1 levels were also significantly higher in the Achilles tendinous tissues of OTR mice and was 3.2-fold higher when compared to the control, while patellar tendon tissues showed a 2.3-fold change compared to control but without statistical significance (Fig 1B, C). These results indicate that only excessive mechanical loading conditions, ITR and OTR, trigger the release of HMGB1 from the tendon cells into the tendon matrix. In order to confirm that the higher HMGB1 concentration in ITR and OTR groups is not due to massive cell destruction during sample preparation, we measured the total DNA content in all samples, and they were equivalent but significantly lower than total lysed tendon samples (Fig 1D). Additionally, the *in vivo* results are supported by *in vitro* cell mechanical stretching experiments; that is, mechanical overloading induces release of HMGB1 from the tendon cells to the culture media (**S2 Fig**).

### Short term ITR induces inflammatory cell infiltration in tendon

To investigate whether intensive mechanical loading induces inflammatory cell infiltration in tendinous tissues, we performed immunostaining of Achilles tendinous tissues after MTR, ITR, and OTR with inflammatory cell marker CD68. CD68 staining was absent in the control and MTR (Fig 2A, B), but was detected in the Achilles tendon sections of mice on the ITR regimen (Fig 2C). The induction of inflammatory cell infiltration by ITR implies that HMGB1 may invoke an inflammatory reaction in tendon (arrow heads, Fig 2C). However, CD68 was found negative in OTR regimen (Fig 2D). Although OTR could induce HMGB1 release within a short period of time (Fig 1A, B), it may not have sustained long enough to induce any inflammatory cell infiltration unless excessive mechanical loading is repeated for a prolonged period.

**Fig 2.**
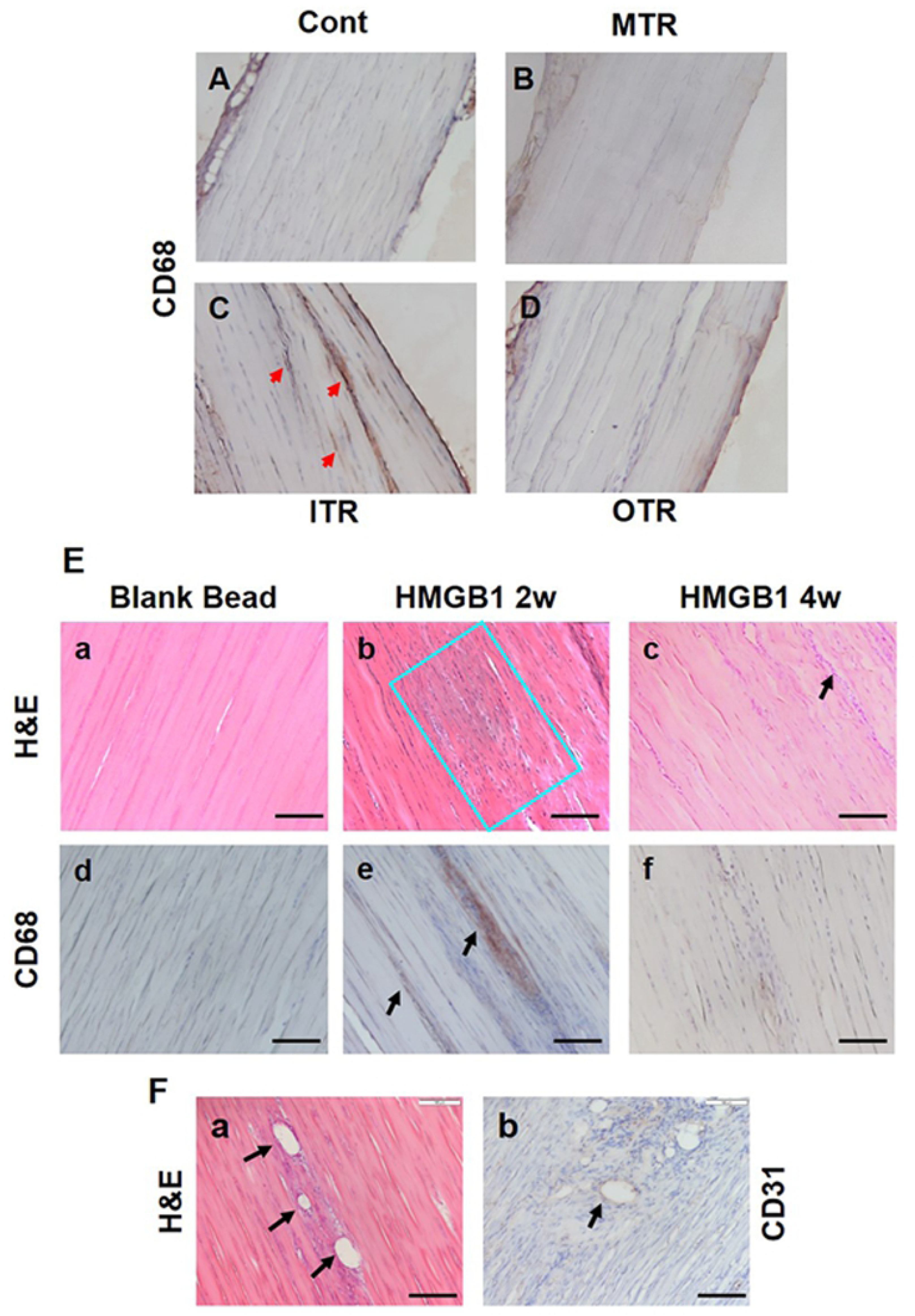
Inflammatory cells infiltrate to tendon matrix under short term ITR, and implanted HMGB1 induces hypercellularity, inflammatory cell infiltration, and angiogenesis in tendon. Mouse Achilles tendons (ATs) are stained with CD68 for inflammatory cells like macrophages and monocytes. **(A)** Mouse AT with cage activity is negative for CD68 stain. **(B)** Similar to cage control group, MTR group shows no CD68 positive staining. (**C)** AT from short term ITR group shows several positively stained regions for CD68 (red arrows). (**D)** CD68 is negative in OTR tendon tissue. The brown signals at the edge of the tendon in adjacent soft tissue are easy to trap antibody that may result in a false negative signal, therefore, signal outside the tendon proper is not considered. **(E**) (**a**) H&E stain of control tendon implanted with blank beads shows no cell proliferation. (**b**) Tendon section with implanted HMGB1 beads shows extensive cell proliferation (highlighted in the blue box) after implantation for 2 weeks. (**c**) The implantation site after 4 weeks; higher number of cells (arrowhead) compared to control can be seen but is much less compared to the 2 weeks group. (**d**) Control with no positive IHC stains for CD68. (**e**) Positive CD68 staining in HMGB1 implanted sample for 2 weeks shows inflammatory cell infiltration (arrows). (**f**) 4 weeks implantation group shows minimal positive staining for CD68. Figures show representative results from at least 3 samples. (**F**) (**a**) Vessel-like structures are present in tendon matrix after implantation of HMGB1 beads for 2 weeks (arrows point to vessels. H&E staining). (**b**) In the same group, extensive angiogenesis (arrow, **b**), as shown by positive IHC staining for CD31, is detected in the tendon matrix. No similar structure was found in control group with blank beads or in the 4 weeks implantation group. Bar: 100 µm.

To confirm the likelihood that HMGB1 causes inflammatory response in tendon, we implanted HMGB1 in alginate beads to rat tendon and immunostained tendon sections for CD68 and CD31 after 2 and 4 weeks. The H&E stain of control tendon implanted with blank beads shows no cell proliferation (Fig 2E, a), while tendon section with implanted HMGB1 beads shows extensive cell proliferation (highlighted in the blue box) after 2 weeks (Fig 2E, b). The implantation site after 4 weeks showed higher number of cells (Fig 2E, c**, arrow**) compared to control, but is much less compared to the 2 weeks group. Control did not show positive IHC stains for CD68 (Fig 2E, d), but positive CD68 staining in HMGB1 implanted sample for 2 weeks showed inflammatory cell infiltration (Fig 2E, e**, arrows**). The four weeks implantation group showed minimal positive staining for CD68 (Fig 2E, f). Moreover, implantation of HMGB1 beads at two weeks resulted in the formation of vessel-like structures (Fig 2F, a, **arrows**). Positive IHC staining for CD31 reveals extensive angiogenesis in the group (Fig 2F, b**, arrow**). Collectively, these data show that under prolonged repetitive mechanical overloading conditions (ITR), HMGB1 is released into the Achilles and patellar tendon matrix, leading to hypercellularity, inflammatory cell infiltration, and angiogenesis in tendon.

### GL blocks HMGB1-induced tendon inflammation *in vivo*

To determine whether GL can negate the inflammatory effects of HMGB1 released to tendon matrix by the short term ITR regimen (3 weeks), we administrated GL to mice by IP injection on the ITR regimen 15 min before they started the treadmill running. Prior to this assay, we determined whether injected GL can be transported and remain in the tendon region after injections. Quantification of GL using thin layer chromatography 3 hrs after injection showed significant levels of GL in mouse patellar and Achilles tendons (**S3 Fig**). GL injection into control mice did not alter the PGE_2_ levels when compared to the mice without injection (Fig 3A). However, PGE_2_ levels were significantly higher in ITR mouse tendons. Specifically, measurement by ELISA showed 1.5 and 1.6-fold increase in AT and PT, respectively, compared to control mouse tendons after an ITR regimen. However, daily GL injection prior to ITR inhibited PGE_2_ production (Fig 3A). Similar effects were observed with MMP-3 levels in mouse tendons after GL injections. MMP-3 levels were significantly elevated in ITR group (1.9 and 1.8-fold increase compared to control), but GL negated the enhanced MMP-3 production (Fig 3B). While statistically not significant, the MMP-3 levels in GL+ITR group appeared to be higher than the control group. Collectively, these results suggest that injection of GL, an inhibitor of HMGB1, reduces inflammation marked by high levels of PGE_2_ in tendon *in vivo*. These results are supported by the *in vitro* data, which showed that exogenous HMGB1 induced high levels of PGE_2_ and MMP-3 production in tendon cells, and GL inhibited the inductions (**S4 Fig**).

**Fig 3.**
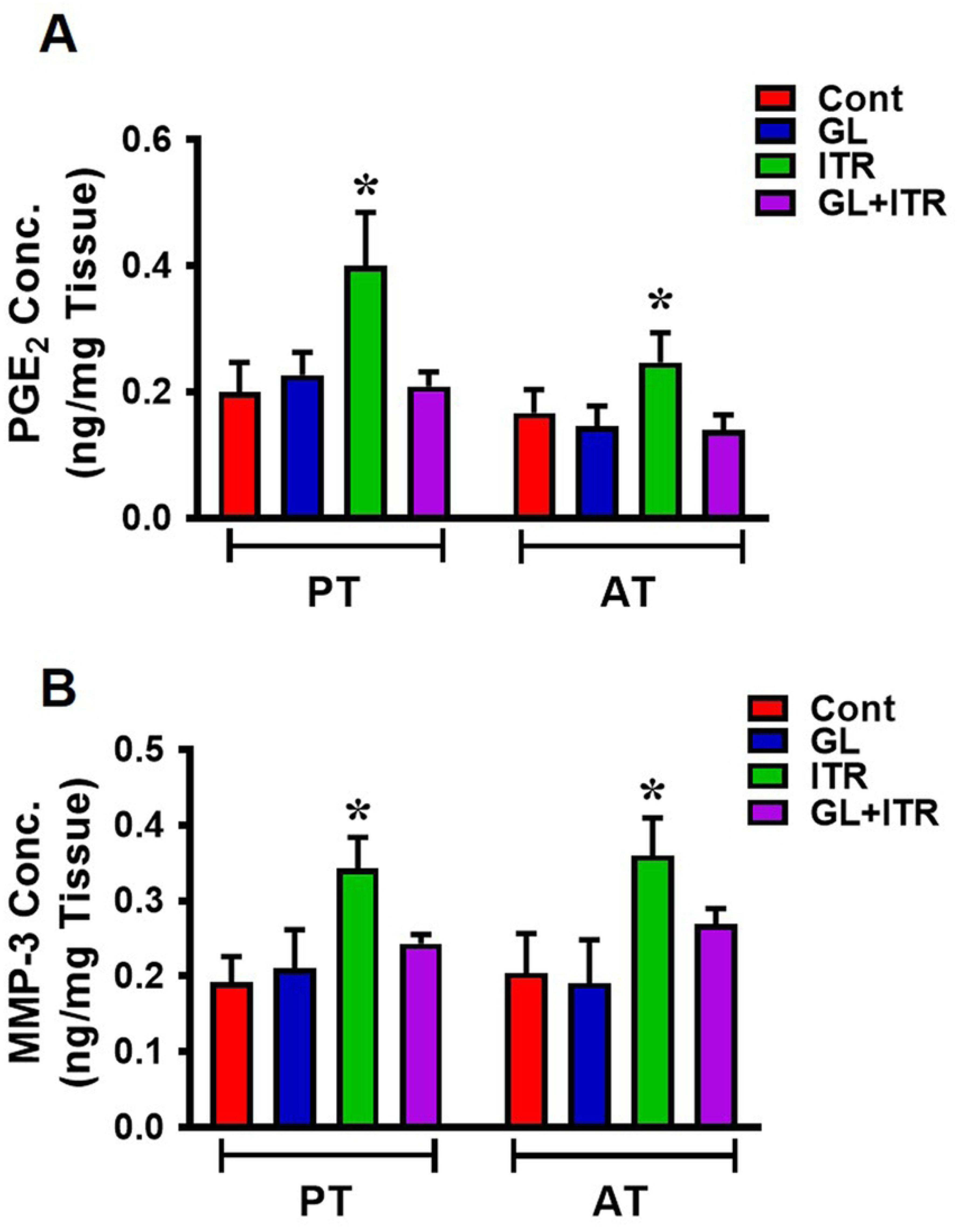
GL injection blocks short term ITR-induced inflammatory reactions in mouse patellar and Achilles tendon tissues *in vivo*. (**A**) PGE_2_ concentrations significantly increase in patellar tendon (PT), and Achilles tendon (AT) after short term (3 weeks) ITR, and GL administration reduces PGE_2_ levels in both tendons. (**B**) Similarly, MMP-3 levels significantly increase in PT and AT in ITR group and GL administration blocks these effects. Data represent mean ± SD. n = 6. \**P* < 0.05.

### Long term-ITR for 12 weeks induces tendinopathy at the tendinous tissue proximal to tendon insertion site

Having established that a short term ITR (3 weeks) induces inflammatory responses in tendon, we next investigated the effect of Long term-ITR (Lt-ITR) on tendinopathy development in mouse Achilles tendon. After 12 weeks of ITR, no obvious structural and compositional changes were found in the middle 1/3 section in Achilles tendon, but the histological analysis at the tendinous tissue near the tendon insertion site revealed typical tendinopathic changes including change in cell shape, accumulation of GAG, and increase in SOX-9 staining (Fig 4A-B). Normal tendon cells in control group are tightly packed in the collagen tissue are largely spindle shaped (Fig 4A, a-c). However, many of the cells in Lt-ITR mouse Achilles tendon were round with lacunae around the cells, which is a typical chondrocyte appearance (Fig 4A, d-f). Semi-quantification of the percentage of round cells near the end site of Achilles tendon showed around 30% of cells are round with cartilage lacunae (Fig 4A, g). Also, there is minimal GAG in the normal tendon (Fig 4A, a-c), while Lt-ITR induced significant GAG accumulation (Fig 4A, d-f). Additionally, there were no round cells or SOX-9 staining in the control (Fig 4B, a, b), while some of those round shaped cells in Lt-ITR mouse tendon were positive for SOX-9 (Fig 4B, c, d). Semi-quantification revealed that about 20% of the cells in 12-week Lt-ITR mouse tendons near the insertion site were positive for SOX-9 (Fig 4B, e).

**Fig 4.**
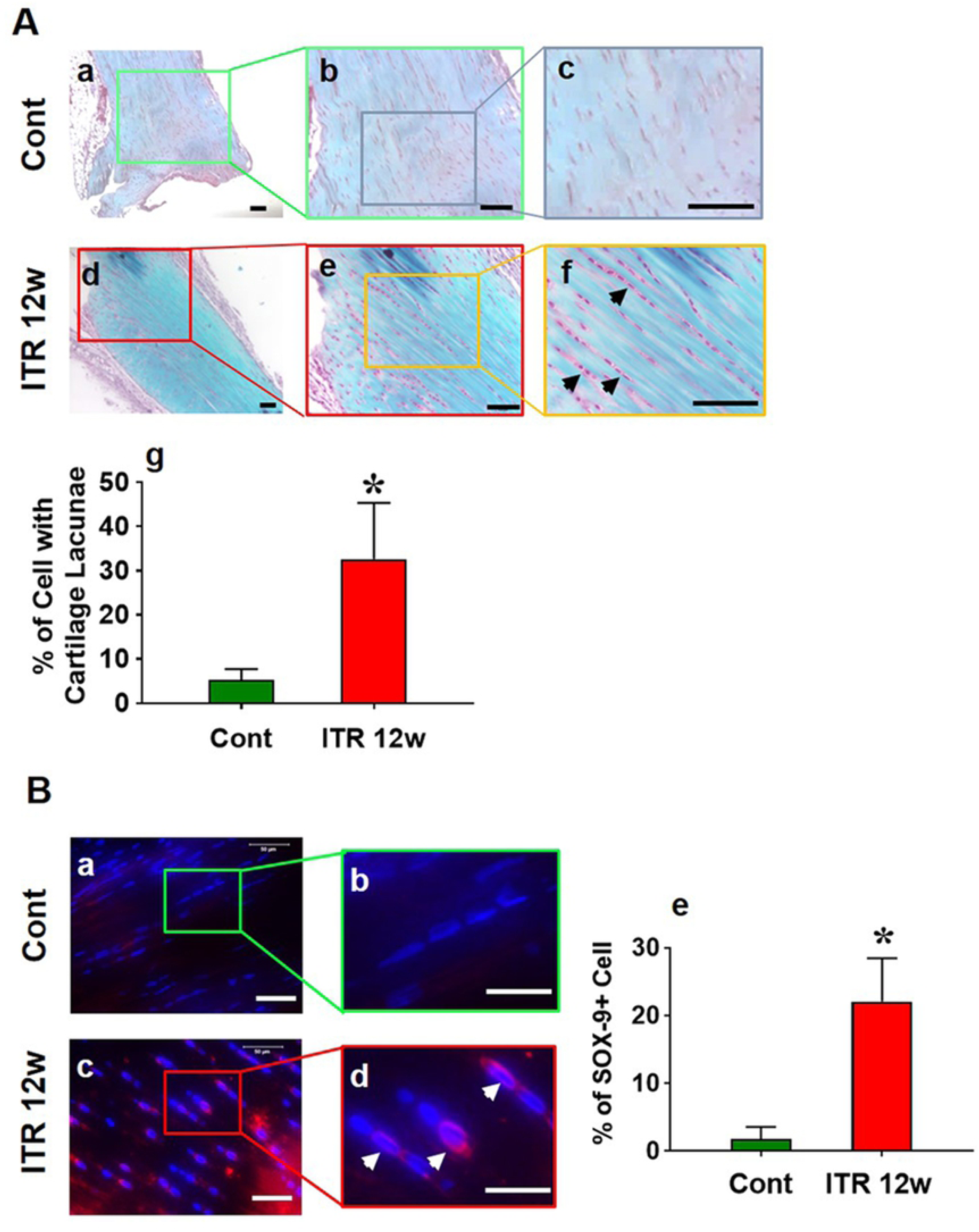
Tendinous tissue near Achilles-bone insertion site shows cell morphology change, GAG deposition, and cartilage marker SOX-9 expression after 12-week Lt-ITR. **(A)** Alcian blue and nuclear fast red staining of tendinous tissue. (**a, b, c**) Tendon from cage control mice shows regular tendon matrix structure and spindle-shaped cell morphology, and tendon cells are tightly packed amongst collagen fibers with little space between the cell body (**a-c**: 4x, 10x, and 20x magnifications). There is also minimal staining for GAG in the control group. In ITR tendon, tendon matrix (**d, e, f**) contains chondrocyte-like “round” cells with cavities called cartilage lacunae (**f, black arrows**), with obvious “blank” area between cells and extracellular matrix (**d-f**: 4x, 10x, and 20x magnifications). Moreover, extensive “blue” staining of GAG is shown. (**g**) Semi-quantification of the percentage of round cells with cavities in a 20x field on the end site of Achilles tendon shows around 30% round cells with cartilage lacunae. (**B**) SOX-9 staining. (**a, b**) Achilles tendons from control group show minimal staining for SOX-9. (**c, d**) Achilles tendons from ITR group show SOX-9 staining in the tendon (white arrows). (**e**) Semi-quantification shows about 20% of total cells are SOX-9 positive in the tendon while no cells in control group express SOX-9. Data represent mean ± SD. n = 4. \**P* < 0.05. Bar: 50 µm.

### Long term-ITR for 24 weeks induces tendinopathy and administration of GL prevents tendinopathy development

The above findings indicate that 12 weeks of Lt-ITR induces the development of insertional Achilles tendinopathy. Therefore, we decided to extend the treadmill running period to 24 weeks to maximize the tendinopathic effects on mice while testing the inhibitory effect of GL on HMGB1 in preventing Achilles tendinopathy. We first checked the presence of HMGB1 in the tendon matrix near the insertion site. HMGB1 was not present in the control or GL only treated group as expected, but HMGB1, as well as CD68, was present in the tendon matrices of the mice after 24 weeks of Lt-ITR (**S5 Fig**). In order to closely evaluate the Lt-ITR effect and GL inhibitory effect, we divided the Achilles tendon near the insertion site into two areas, the proximal region (∼300 µm from the end of tendon tissue), which belongs to Achilles tendinous tissue (Figs 5, 6, yellow boxes), and the distal region (Figs 5, 6, green boxes) next to the tendon-bone insertion, which is very near the end of the tendon tissue that is considered as part of transitional zone between tendon and heel bone. We found that a small number of chondrocyte-like cells exist in the distal region in the control group. With this in mind, we focused on the proximal region since it represents the site of degenerative changes in tendon rather than the region of possible pre-existing chondrocyte-like cells.

**Fig 5.**
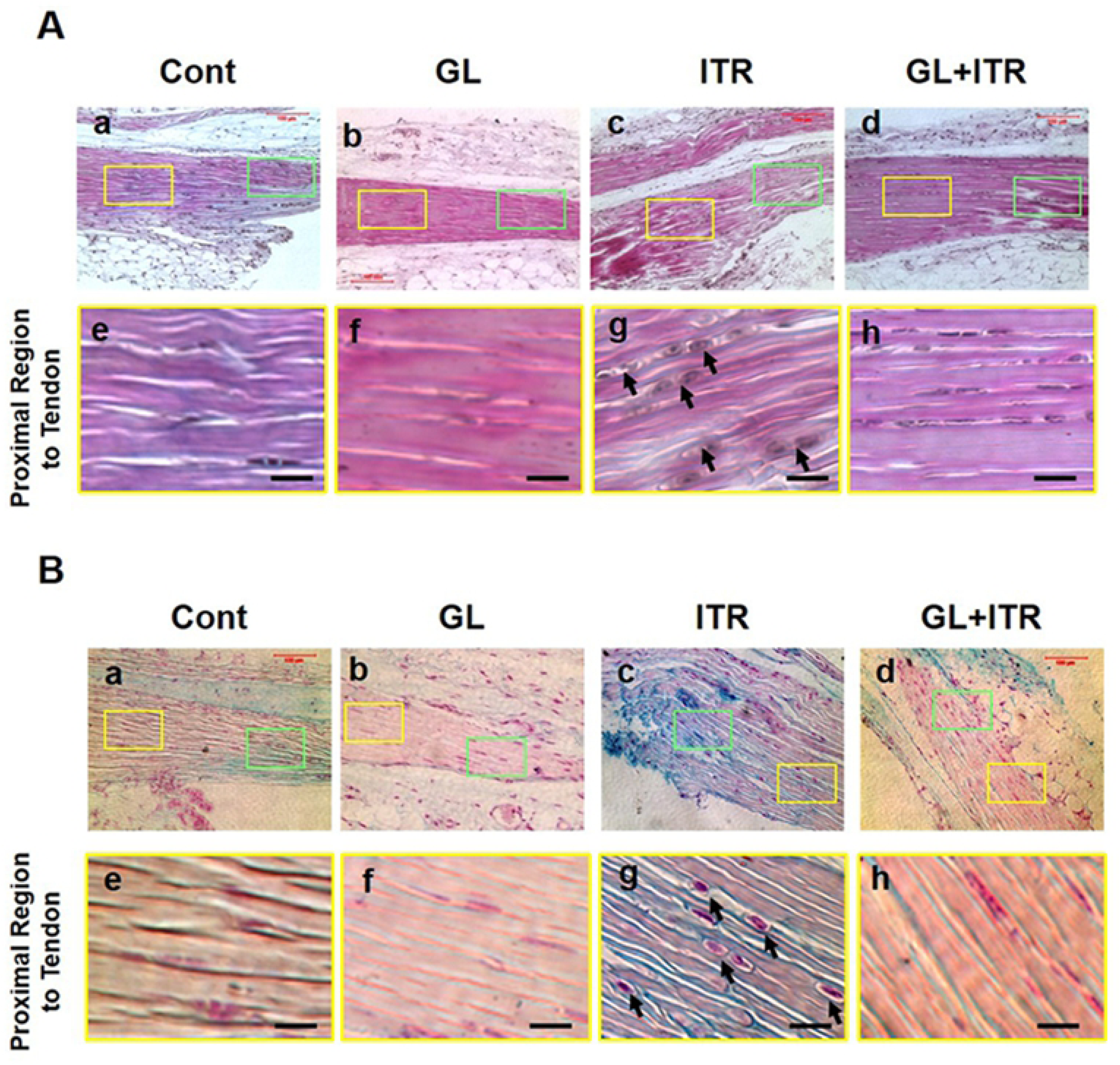
GL attenuates the transformation of tendon cells into chondrocyte-like cells and prevents GAG deposition induced by 24-week Lt-ITR in tendinous tissue near the insertion site of mouse Achilles tendon. (**A**) Achilles tendon in (**a**) Cage control (Cont). (**b**) GL injection only (GL). (**c**) Intensive treadmill running (ITR). (**d**) GL injection+ Intensive treadmill running (GL+ITR). Also, the yellow boxes in (**a-d**) indicate the proximal region of Achilles tendon, which is away from the Achilles tendon-bone insertion site, whereas green boxes point to the area nearby where the Achilles tendon-bone insertion site is located. We focus on the tendinous region indicated by yellow boxes. (**e, f**) Control (Cont) and GL only groups do not show round cells with cavities, meaning no cartilage cells exist in this tendinous region. (**g**) Tendon at the proximal region (yellow) in ITR group contains numerous chondrocyte-like cells with cavities (black arrows). (**h**) GL treatment of ITR group results in the presence of few chondrocyte-like cells in the proximal region of Achilles tendon, indicating that GL treatment attenuates chondrocyte-like cell differentiation induced by ITR. (**B**) Alcian Blue staining shows the overall GAG deposition in the same groups as above. (**e, f**) In Cont and GL injection alone groups, minimal staining of GAG is present in the proximal region of Achilles tendon (yellow boxes). (**g**) In ITR group, strong staining of GAG in the proximal region is shown, with chondrocytes-like cells present in the matrix (black arrows point to cavities around the cells called cartilage lacunae). Extensive GAG staining is also evident in the distal region (**c**, green box), which is expected because this region is considered to be a part of fibrocartilage transitional zone between Achilles tendon and heel bone. (**h**) In GL treated ITR group, there is minimal staining of GAG in the proximal region of tendinous tissue. Bar: 25 µm.

**Fig 6.**
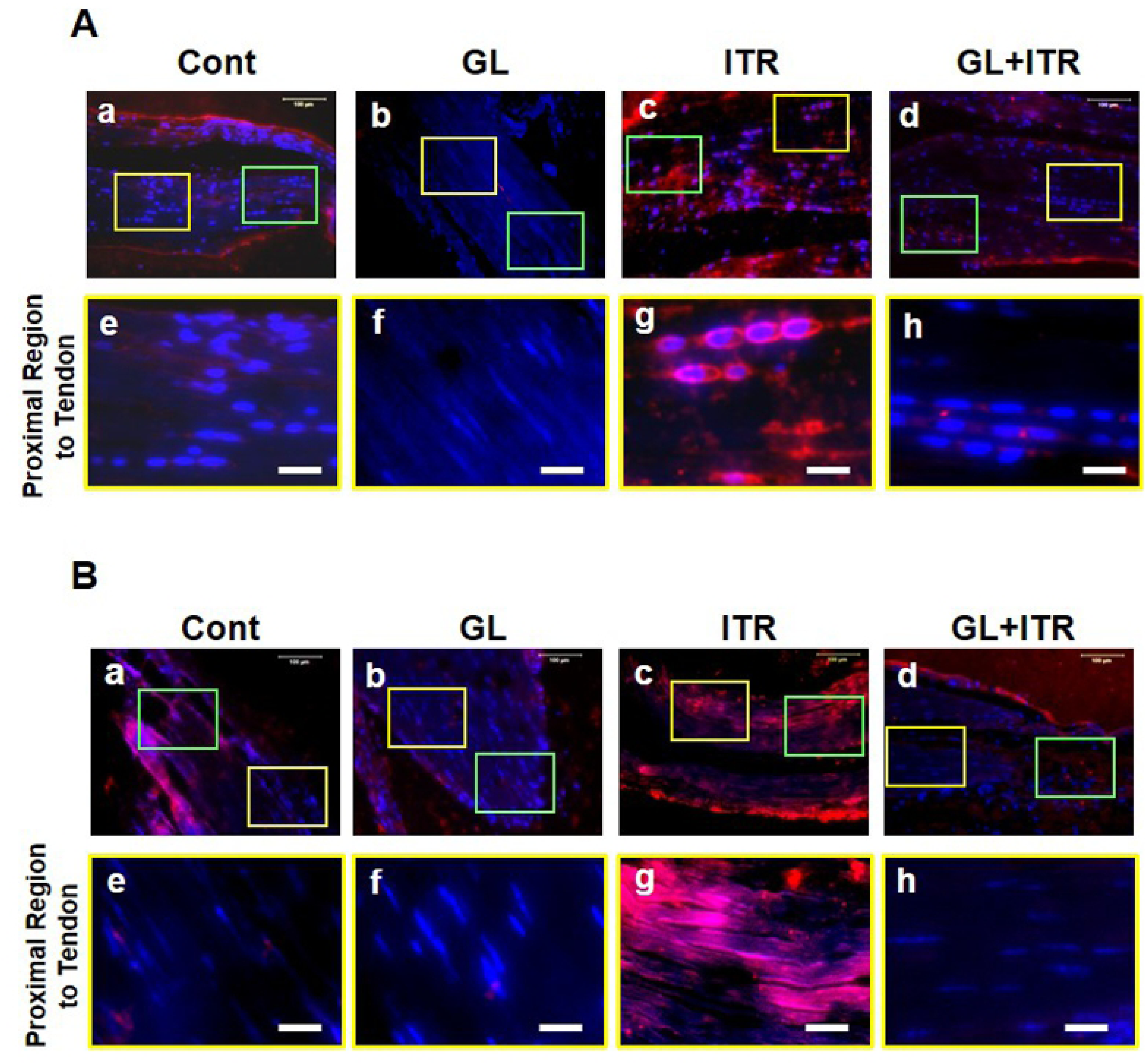
GL treatment reduces the expression of SOX-9 and deposition of collagen II induced by 24-week Lt-ITR in tendinous tissue near the insertion site of mouse Achilles tendon. (**A**) SOX-9 staining in cage control group (**a**), GL injection alone group (**b**), ITR group (**c**), and GL+ITR group (**d**). Yellow boxes indicate the proximal region of Achilles tendon-away from insertion site, whereas green boxes point to the distal region closer to the insertion site. Since the distal region is part of the fibrocartilage transition zone, our analysis is only focused on the proximal region of the tendinous tissue. (**e**) Proximal region of the control group shows round cells without SOX-9 staining. (**f**) No SOX-9 staining is detected in GL group at the proximal region. (**g**) Strong SOX-9 staining, along with the round shaped cells, are shown in ITR group. (**h**) There is only minimal SOX-9 signal in the same proximal region of Achilles tendon in the GL+ITR group. (**B**). Collagen II staining in the same groups as above. Collagen II is mostly negative in the control and GL groups (**e, f**), but the staining of collagen II is extensive in the proximal region of Achilles tendon in the ITR group (**g**). However, after GL treatment (GL+ITR), ITR-induced collagen II expression in the proximal region of Achilles tendon is minimal (**h**). Bar: 25 µm.

We found that after 24 weeks of Lt-ITR, the proximal site of Achilles tendon contained cells with round shape (arrowheads in Fig 5A, g), compared to more elongated cells in the cage control and GL treatment group alone groups (Fig 5A, e, f). However, in Lt-ITR mice treated with daily injection of GL, no change in cell shape was observed (Fig 5A, h). In addition, in these Lt-ITR mice, extensive GAG staining in the tendon was present. Such GAG accumulation was prevented by GL administration in the group (Fig 5B**, bottom panel**). Also, Lt-ITR induced the expression of chondrogenic markers (SOX-9 and Col II) in Achilles tendon, and GL inhibited the expression of SOX-9 and Col II confirmed by immunofluorescence staining (Fig 6A, B**, bottom panel**). Collectively, these findings suggest that Lt-ITR for 24 weeks induces degenerative changes, typical of insertional tendinopathy at the proximal site of Achilles tendon, and that injections of GL blocks the tendon’s degenerative changes due to mechanical overloading on the tendon.

## Discussion

Tendinopathy affects large populations in both athletic and occupational settings. Management of this tendon disorder is an ongoing challenge due to lack of understanding of the precise molecular mechanisms underlying the development of tendinopathy (44, 45). Inflammation is thought to be a major contributor to the development of tendinopathy (11, 46). Previously, we have shown that excessive mechanical loading induces inflammatory mediator PGE_2_ in tendon cells and tissues (15, 17). A few other studies have shown that upregulation of HMGB1 is associated with shoulder tendon injury/tendinopathy in patients and indicated that it would be a valuable target for tendinopathy management (37–39, 47). In this study, we extend these investigations to examine the role of HMGB1, an inflammatory alarmin molecule, in tendinopathy development due to mechanical overloading conditions. We show that HMGB1 is released to the extracellular space of tendon cells under mechanical overloading conditions thereby eliciting the cells’ inflammatory and catabolic responses marked by elevated PGE_2_, and MMP-3 production in a rodent model. Moreover, we show that by daily IP injection, GL reduces the inflammatory/catabolic reactions marked by high levels of production in PGE_2_ and MMP-3, in overloaded mouse tendons *in vivo*. Finally, GL administration in mice that underwent long term intensive treadmill running blocks the development of degenerative tendinopathy characterized by the presence of chondrocyte-like cells, accumulation of proteoglycans, chondrogenic marker SOX-9 expression, and high levels of collagen type II production.

Although the presence of inflammation in tendinopathic tendons is highly debated, the contribution of immune cells and inflammatory mediators to tendinopathy development are increasingly recognized. Recent investigations in human tissues and cells from tendinopathic patients strongly support that inflammation is involved in tendinopathy (11–13). Infiltration of inflammatory cells like macrophages and mast cells has been reported in early supraspinatus tendinopathy patients (11). Tendinopathy may not progress through a classic inflammatory pathway but may rather involve a local sterile inflammation initiated by overloaded and damaged cells that could release molecules functioning as danger signals. Alarmins including HMGB1 are implicated as key effectors in the activation of immune system that may be important in the pathogenesis of tendinopathy (36). However, there is limited data regarding the potential role of HMGB1 in tendinopathy development. By using *in vivo* and *in vitro* models, we show for the first time that HMGB1 induces inflammatory reactions in tendon cells and tendon matrix, a hallmark of early stages of tendinopathy, and injection of HMGB1 inhibitor, GL, abolishes the development of degenerative tendinopathy.

Recent findings with clinical samples of tendinopathy indicate that HMGB1 is present and likely plays an important role in driving early stages of tendinopathy (11). The tissue and cells derived from tendinopathic and ruptured Achilles tendons show evidence of chronic (non-resolving) inflammation (13). Additional clinical studies support this finding showing enhanced levels of HMGB1 in early stage supraspinatus tendinopathy tissues compared to normal tissues and late stage tendinopathy tissues (37, 38). Moreover, the *in vitro* study shows that recombinant HMGB1 induces significant inflammatory mediators such as IL-1β, IL-6, IL-33, CCL2, and CXCL-12 (37). The findings of this study further links HMGB1 with the inflammatory responses induced by mechanical overloading of tendon to the developmental course of tendinopathy. In other organs, once released following trauma or severe cellular stress thereby triggering sterile inflammation in injured tissues, HMGB1 is implicated as a causative factor in many diseases, *i.e.* sepsis, rheumatic arthritis, pancreatitis, ischemia-reperfusion injury, and gastrointestinal disorders (34, 48). Inhibiting HMGB1 using anti-HMGB1 neutralizing antibody attenuated the development of pancreatitis and associated organ dysfunction (49). Blocking HMGB1 activity is therapeutic in arthritis, because administration of either anti-HMGB1 or A-box of HMGB1 in collagen type II-induced arthritis significantly attenuated the severity of disease (50). Thus, HMGB1 may represent a new target of therapy of inflammation-related diseases such as tendinopathy. This is supported by the finding that use of GL, a specific inhibitor of HMGB1, suppresses inflammatory responses as defined by PGE_2_ in tendon cells and prevents tendinopathy development.

In this study, HMGB1 was shown to be released into tendon matrix in response to mechanical overloading conditions (ITR). But the exact “release modes” are not clear. The HMGB1 may be actively released by tendon cells due to excessive mechanical stress on the cells, as suggested by *in vitro* data of this study; or it may be from passive release by loading-induced cellular necrosis. Additionally, macrophage/monocytes recruited to tendon tissue during overloading may also release HMGB1. All “release modes,” would lead to inflammatory responses in tendon matrix.

It is known that HMGB1 acts as a chemoattractant in various cell types like macrophages, neutrophils, mesoangioblasts, and osteoclasts (33, 34). Our study also demonstrates this well-known property of HMGB1 in tendons. After initial tendon microinjury by repetitive mechanical over loading such as long-term intensive treadmill running in this study, inflammation can occur with influx of white blood cells, and HMGB1 by its chemoattractant property may recruit neutrophils, monocytes, and macrophages to sites of injury. While exploring the HMGB1 effect *in vitro*, we found that this mediator did promote tendon cell migration and inflammatory reactions but did not induce their proliferation (data not shown). Interestingly, in the HMGB1 implantation experiment, HMGB1 induced hypercellularity in tendon tissues (Fig 2E). It is likely that HMGB1 exerts its function by recruiting inflammatory cells to the “injury site”, and then initiates the release of cytokines (e.g. IL-6, IL-1β, IL-6, and IL-8) from the inflammatory cells. This should be investigated in future studies.

In this study, we used GL to inhibit HMGB1-induced inflammation as defined by PGE_2_ in contrast to inflammatory markers such as IL-1β, IL-6, and IL-8, since clinically, inflammation reduction is achieved by using non-steroidal anti-inflammatory drugs (NSAIDs), which inhibit COX and as a result, reduce PGE_2_. Moreover, the amplification of the PGE_2_ biosynthesis pathway by HMGB1/IL-1β is suggested as an important pathogenic mechanism perpetuating inflammatory and destructive activities in rheumatoid arthritis (51).While it is currently unclear whether our findings will reflect the actual mechanisms of tendinopathy development and treatment in humans, recent studies demonstrated that HMGB1 is present in human tendinopathic tendons and regulates cellular inflammation and protein production *in vitro* (37, 38). This suggests that HMGB1 plays a similar role in the development of tendinopathy due to mechanical overloading.

The specific inhibitor of HMGB1, glycyrrhizin (GL), is a natural glyco-conjugated triterpene present in licorice plant. It blocks prostaglandin production and inflammation (52). Topically, it has been in use for the treatment of tendinitis, bursitis, and gum inflammation. It has been used in preclinical investigations to inhibit HMGB1 signaling to treat inflammation in lung and liver diseases (42). GL has a long history of well-known anti-inflammatory effects (53–55), and studies show that it can inhibit the chemoattractant and mitogenic activity of HMGB1 by direct binding (52, 56). It has also been administered to patients with hepatitis B and C and is considered to be safe for human consumption (52, 57, 58), but GL may have off-target effects other than inhibition of HMGB1. The side effect of GL is mainly from its metabolic product glycyrrhetic acid after oral ingestion and catalyzed by bacteria in gut(59), which is irrelevant to IP injection of GL that we used in this study. Therefore, GL is safe for *in vivo* use and has minimal off-target effects when IP injection is used for GL delivery.

Based on the findings in this study, we propose a pathological tendinopathy model focusing on the role of HMGB1 in tendinopathy development and the subsequent degenerative changes. Mechanical overloading of tendon results in micro-tears of tendon matrix and/or tendon cells, and as a result HMGB1 is released from stressed or injured tendon cells. The extracellularly released HMGB1 attracts inflammatory cells (e.g. macrophages) to the injury site, and they release inflammatory cytokines. The resident tendon cells are also activated and shift to a pro-inflammatory phenotype. The proliferation of tenocytes, ingrowth of blood vessels, and destruction of well-organized collagen matrix, result in compromised mechanical property of the tendon that is vulnerable to even normal mechanical loading. Due to the persistence of the overloading, as opposed to one-time or modest loading, the inflammation status does not get resolved but rather gets amplified. As a consequence, chronic sterile inflammation persists in tendon tissue, which leads to degenerative changes that eventually lead to the development of full-blown tendinopathy. Our previous studies showed that PGE_2_ treatment induced non-tenogenic differentiation of tendon stem cells into adipocyte, chondrocyte, and osteocytes both *in vivo* and *in vitro* (17, 60), and these studies help to explain how chronic inflammation may result in a chondrogenic phenotype change in our treadmill running overloading model.

In conclusion, our results support that HMGB1 released to tendon matrix due to mechanical overloading induces tendinopathy development by initiation of tendon inflammation and eventual tendon degeneration. These results provide evidence for the role of HMGB1 as a therapeutic target to prevent tendinopathy before its onset and block further development at its early inflammation stages. The inhibition of tendinopathy development by GL administration in this study also suggests that GL may be used as a therapeutic agent to prevent tendinopathy development.

## Supporting information

**S1 Fig. HMGB1 is present in mouse tendon and located in the nuclei of tendon cells without mechanical overloading. (A)** A standard Western blot shows the presence of HMGB1 in both tissues (two samples from different animals) and cells (from two different wells) of patellar tendon (PT) and Achilles tendon (AT). Total protein was extracted from rat Achilles and patellar tendons and cells using T-PER buffer. After quantification, 20 µg of total protein from each tendon sample was separated on a 10% SDS-PAGE, transferred onto a nylon membrane and incubated with rabbit anti-HMGB1 primary antibody (rabbit anti-mouse, 1 µg/ml, Cat # ab18256, Abcam) followed by goat anti-rabbit infrared tag conjugated secondary antibody (1:5,000 dilution, Cat # C30409-07, LI-COR Biosciences, Lincoln, NE) following the manufacturer’s instructions. Positive signals were detected via the Odyssey CLx infrared imaging system (LI-COR Biosciences, Lincoln, NE). β-actin served as internal control. **(B)** Immunostaining of tendon tissue stained for HMGB1 without penetration with detergent shows that HMGB1 is minimal in tendon matrix. **(C)** HMGB1 staining in the tendon with Triton X-100 penetration treatment shows that HMGB1 is located in tendon cell nucleus and cytoplasm. **(D)** HMGB1 staining of tendon cells in culture. Most cells contain HMGB1 in their nuclei (red). (**E**) Hoechst H33342 stained nuclei (blue). **(F)** Overlay of both staining **(D, E).** While HMGB1 is located in the nuclei of most cells (pink), it is missing in some cells. Bar: 50 µm.

**S2 Fig. Mechanical overloading of tendon cells *in vitro* induces release of HMGB1 to culture media. (A) (a, d)** Unstretched control cell nuclei stained positive for HMGB1 (pink). (**b, e**) 4% stretched cell nuclei also stained positive for HMGB1. (**c, f**) 8% stretched cells show that the majority of cells lose HMGB1 in their nuclei, indicating that their cells have released HMGB1 to culture media under 8% mechanical overloading. Semi-quantification analysis confirms the results (**g**). Specifically, without mechanical loading or 4% stretching, more than 95% of tendon cells are stained positive for HMGB1. In contrast, there is only about 35% cells that are positive staining with HMGB1, which represents 65% reduction in HMGB1 positive nuclei due to mechanical overloading on the tendon cells. **(B)** The levels of HMGB1 in culture media were measured using ELISA kits. It is shown that 8% stretch significantly increases HMGB1 levels compared to control and 4% stretch. The cell stretching experiments were done according to our published protocol (15, 61). All data are means ± SD. n = 6. **\****P* < 0.05. Bar: 50 µm.

**S3 Fig. IP injection results in the presence of GL in tendon.** Three hours after IP injection, significant amounts of GL, quantified using thin layer chromatography (62), are detected in mouse tendons. Amount of GL is minimal in mouse tendons without IP injection of GL, but there is 13-fold increase of GL in PT and 6.8-fold increase in AT. PT – patellar tendon, and AT – Achilles tendon. n = 4. *\*P* < 0.05.

**S4 Fig. GL blocks PGE_2_ and MMP-3 production induced by HMGB1 in tendon cells.** (**A**) Tendon cells derived from rat Achilles tendons were treated with 10 µg/ml HMGB1 or 10 µg/ml HMGB1+ 200 µM GL in culture, 10 ng/ml IL-1β served as a positive control. PGE_2_ levels determined by ELISA, significantly increase at 10 µg/ml HMGB1 treatment at 0.5, 2, and 4 hrs, and combined treatment with GL (200 µM) mitigates the effects of HMGB1. (**B**) HMGB1 treatment (10 µg/ml) of tendon cells significantly increase the production of MMP-3 (ELISA quantification) by tendon cells in culture medium, but addition of 200 µM GL with HMGB1 reduces MMP-3 to a similar level as the non-treated control. Data represent mean ± SD. n = 4. \**P* < 0.05.

**S5 Fig. HMGB1 is present near the insertion site of Achilles tendon after 24 weeks ITR. (A, B**) In the proximal region of tendinous tissue near the mouse Achilles tendon-bone insertion site, HMGB1 staining is minimal in control and GL only groups. **(C)** HMGB1 is present in tendon matrix in the treadmill running group (arrows). **(D)** HMGB1 is also detected in the tendon matrix of GL+ITR group (arrows). **(E, F)** CD68 staining is negative in cage control and GL injection only group (yellow arrows). **(G)** CD68 is positive in ITR group (arrows) and gathered in a clustered form. **(H)** No positive CD68 signal in the GL-treated ITR tendon tissue. Bar: 50 µm.

## Acknowledgements

We thank Dr. Bhavani P Thampatty and Dr. Huiyan Wu for their assistance in the preparation of this manuscript.

